# Genome-wide association mapping of correlated traits in cassava: dry matter and total carotenoid content

**DOI:** 10.1101/102665

**Authors:** Ismail Y. Rabbi, Lovina I. Udoh, Marnin Wolfe, Elizabeth Y. Parkes, Melaku A. Gedil, Alfred Dixon, Punna Ramu, Jean-Luc Jannink, Peter Kulakow

## Abstract

Cassava (*Manihot esculenta* (L.) Crantz) is a starchy root crop cultivated in the tropics for fresh consumption and commercial processing. Dry matter content and micronutrient density, particularly of provitamin A – traits that are negatively correlated – are among the primary selection objectives in cassava breeding. This study aimed at identifying genetic markers associated with these traits and uncovering the potential underlying cause of their negative correlation – whether linkage and/or pleiotropy. A genome-wide association mapping using 672 clones genotyped at 72,279 SNP loci was carried out. Root yellowness was used indirectly to assess variation in carotenoid content. Two major loci for root yellowness was identified on chromosome 1 at positions 24.1 and 30.5 Mbp. A single locus for dry matter content that co-located with the 24.1 Mbp peak for carotenoid content was identified. Haplotypes at these loci explained a large proportion of the phenotypic variability. Evidence of mega-base-scale linkage disequilibrium around the major loci of the two traits and detection of the major dry matter locus in independent analysis for the white- and yellow-root subpopulations suggests that physical linkage rather that pleiotropy is more likely to be the cause of the negative correlation between the target traits. Moreover, candidate genes for carotenoid (*phytoene synthase*) and starch biosynthesis (*UDP-glucose pyrophosphorylase* and *sucrose synthase*) occurred in the vicinity of the identified locus at 24.1 Mbp. These findings elucidate on the genetic architecture of carotenoids and dry matter in cassava and provides an opportunity to accelerate genetic improvement of these traits.

**CORE IDEAS:**

- Cassava, a starchy root crop, is a major source of dietary calories in the tropics.
- Most varieties consumed are poor in micronutrients, including pro-vitamin A.
- These two traits are governed by few major loci on chromosome one.
- Genetic linkage, rather than pleiotropy, is the most likely cause of their negative correlation.

## INTRODUCTION

Cassava (*Manihot esculenta* (L.) Crantz) is one of the most important food and feed crops in the tropics and Africa accounts for more than half of the total world-wide production of 270.3 million tonnes (http://faostat3.fao.org/, accessed 26.03.2016). Because of its remarkable tolerance to drought (El-Sharkawy, 1993), its ability to grow in poor soils (Cock, 1982), and its perennial nature which allows it to be harvested as and when required, this heterozygous and clonally propagated species plays a particularly important role in food security for millions of small-holder farmers in developing countries. Moreover, cassava is increasingly being cultivated for commercial processing to convert its storage roots into dehydrated chips, flour and starch (Balagopalan, 2002). Dry matter content, of which a large proportion is starch, is therefore a primary factor that defines adoption of new cassava varieties by farmers and the market value of harvested roots (Okechukwu and Dixon, 2008). As a result, breeding of improved varieties with high dry matter content is one of the primary objectives of cassava genetic improvement programs in the world.

Another important target trait for cassava improvement in developing countries is biofortification for micronutrients (Pfeiffer and McClafferty, 2007; Saltzman et al., 2013). Most varieties grown and consumed throughout the world have white storage roots with negligible amounts of micronutrients in general, and provitamin A in particular (Welsch et al., 2010). Dietary diversification and breeding of farmer-preferred improved varieties with higher nutritional density are complementary approaches used in addressing potential micronutrient deficiency associated with consumption of cassava as the sole staple food (Sayre et al., 2011). The crop’s gene-pool exhibits considerable natural variation for storage root carotenoids that can be tapped for breeding of biofortified varieties, with some breeding populations reported to accumulate as much as 25.8 μg/g fresh root weight (Ceballos et al., 2013; Sánchez et al., 2014).

Despite availability of natural genetic diversity in the global germplasm that is relevant to breeding for increased dry matter and total carotenoid contents, improving these traits through phenotype-based recurrent selections is a lengthy process, due to the breeding complexities associated with the species including an annual cropping cycle of 12 to 24 months and low multiplication rate of planting materials. Understanding the genetic basis of variation in these traits is essential for increasing their selection efficiency and the rate of genetic gain. More importantly, several studies using diverse germplasm have reported that dry matter and carotenoid content are negatively correlated, with *r* values ranging from -0.1 to -0.5 (Marín Colorado et al., 2009; Akinwale et al., 2010; Esuma et al., 2012; Ceballos et al., 2013; Njoku et al., 2015). Despite its significant implication in breeding, the genetic basis of this correlation – whether it is due to genetic linkage or pleiotropy – is not understood.

Several mapping studies using either Bulk Segregant Analysis (BSA) or Quantitative Trait Loci (QTL) mapping of S1 or F1 populations have been reported separately for dry matter and carotenoid content (Balyejusa Kizito et al., 2007; Marín Colorado et al., 2009; Welsch et al., 2010; Morillo C et al., 2013; Njoku et al., 2014). The mapping resolution from single-cross experimental populations is expected to be limited due to the use of sparse genetic maps and the limited number of recombination events observed (Hamblin et al., 2011). Moreover, QTLs from such bi-parental populations may not provide insight into the tremendous genetic and phenotypic variation of the larger gene pool (Zhao et al., 2011). The increased availability of genomic resources for cassava, including the chromosome-scale reference genome and integrated linkage map (Prochnik et al., 2012; International Cassava Genetic Map Consortium (ICGMC), 2014) and high-density genotyping using next-generation sequencing (Rabbi et al., 2014a; b) makes it possible to use genome-wide association (GWAS) mapping to dissect the phenotypic diversity of cassava germplasm with respect to dry matter and carotenoid content. GWAS, which takes advantage of natural linkage disequilibrium (LD) generated by ancestral mutation, drift, and recombination events in diverse germplasm, offers the possibility to overcome the shortcomings of traditional bi-parental QTL mapping. These advantages mean GWAS is able to reveal a broader spectrum of trait-linked allelic variation and thus may provide the most useful markers for marker-assisted selection (MAS). Indeed, GWAS has already been applied in other crops such as maize to study the genetic architecture of carotenoid accumulation (Harjes et al., 2008; Owens et al., 2014; Suwarno et al., 2015). In cassava, Esuma et al., (2016) carried out a GWAS study using a panel of partial inbreds (S1 and S2 generation) produced from eight clones. Using this limited number of parents, they reported a single genomic region on Chromosome 1 that underlies the variation in total carotenoid content. However, no joint association analysis examining carotenoids and dry matter content has hitherto been reported. Here, we present the results of a GWAS using a collection of more than 650 cassava clones representing diverse African germplasm genotyped at high-density using genotyping-by-sequencing (Elshire et al., 2011). The population was phenotyped in two locations for three consecutive field seasons. The results of this study will be used to develop efficient strategies to breed for high dry matter and provitamin A content varieties.

## METHODS

### Germplasm

The present work was carried out using the Tropical Manihot Selection (TMS) cultivars developed at the International Institute of Tropical Agriculture (IITA) in Nigeria. This population, also known as the Genetic Gain collection, consist of more than 650 advanced breeding lines and key landraces selected over four decades from 1970 (Okechukwu and Dixon, 2008; Ly et al., 2013). The pedigree of the collection is mainly composed of crosses between germplasm from West Africa and early introductions of CMD-tolerant lines arising from interspecific hybridization between *Manihot glaziovii* and cultivated cassava at the Amani station in Tanzania (Hahn et al., 1980). The collection also includes hybrid germplasm from Latin America (Wolfe et al., 2016).

### Locations and experimental design

The Genetic Gain population was planted using an incomplete block design with two checks per block and single row of either 5 or 10 plants spaced at 1m^2^. Data used for this study was collected in the 2012-2013, 2013-2014, and 2014-2015 field seasons in Ibadan (7.40° N, 3.90° E) and Ubiaja (6.66° N, 6.38° E). The trials are usually planted in June, at the onset of the raining season in South West Nigeria and harvested in June of the following year.

### Assessment of dry matter content and yellow color intensity of storage roots

Dry matter content was assessed using the oven-drying method. Eight fully developed roots were randomly selected from each plot, peeled, chipped and thoroughly mixed. For each sample, 100g was weighed and oven-dried for 48 hours at 104^o^C till constant weight was achieved. The samples were then re-weighed and the dry matter content was expressed as the percentage of dry weight relative to fresh weight.

Because of the well-established linear relationship between intensity of yellow color and carotenoid content in cassava storage roots (Pearson’s coefficient, *r*, ranges from 0.81 to 0.89), we used root yellowness as an indirect measure of carotenoid content (Iglesias et al., 1997; Chávez et al., 2005; Marín Colorado et al., 2009; Akinwale et al., 2010; Sánchez et al., 2014). The relative difference among clones in the Genetic Gain population was assessed using two complementary methods. The first was a visual gradation of yellow color using a standard color-chart starting from one (white) to 7 (deep yellow). Due to the potential subjectivity inherent in visual color scores, we complemented the color-chart method through the use of a Minolta CR-410^®^ chromameter. Approximately 100g of grated samples from freshly peeled roots were placed in transparent Nasco Whirl-Pak^®^ sampling bags and four chromameter measurements taken in different sections of the bag. We chose the commission internationale de l'éclairage (CIELAB) method that records color values in a three-dimensional color space, where the L* coordinate corresponds to a lightness coordinate, and the a* coordinate corresponds either to red (positive values) or to green (negative values). Of importance to this study was the b* coordinate, whose positive values represents yellow while the negative values represent blue. The illuminant used was D65 and calibration was done each day with a white ceramic.

### SNP genotyping

DNA was extracted as described in Rabbi et al. (2014) and Genotyping-by-sequencing was carried out as described by (Elshire et al., 2011). DNAs from the Genetic Gain individuals were digested individually with *ApeK*I, a methylation sensitive restriction enzyme that recognizes a five base-pair sequence (GCWGC, where W is either A or T). The GBS sequencing libraries consisting of 95-plex DNA samples each were prepared by ligating the digested DNA to unique sample identifier barcodes (nucleotide adapters) followed by standard PCR. Sequencing was performed using Illumina HiSeq2500. The sequenced reads from different genotypes were de-convoluted using their unique barcodes and aligned to version 6.0 of the cassava reference genome (www.phytozome.org/cassava) with the Bowtie 2 (Langmead and Salzberg, 2012). SNPs were discovered using the GBS pipeline Version 2 implemented in TASSEL software (Glaubitz et al., 2014) and converted to dosage format (0 = homozygous reference, 1 = heterozygous, 2 = homozygous non-reference alleles). Missing data were filtered as described in (Wolfe et al., 2016) and imputed with the glmnet algorithm in R (http://cran.r-project.org/web/packages/glmnet/index.html) (Wong et al., 2014).

### Phenotypic data analysis

The phenotypic data across two locations and three years was collapsed to single best linear unbiased predictor (BLUP) values for each clone by fitting the following mixed linear model with the *lme4* package in R:

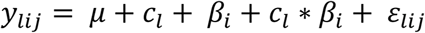

Here, *y*_*lij*_represents raw phenotypic observations, *μ* is the grand mean, *c_l_* is a random effects term for clone with 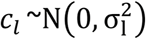, *β_i_* is a fixed effect for the combination of location and year harvested year harvested, *c_l_* * *β_i_* is a random effect for genotype-by-environment variance, and *ε_lij_* is the residual variance, assumed to be random and distributed 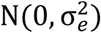. Broad-sense heritability for dry matter content and yellow color intensity was calculated according to (Ly et al., 2013). Genetic correlation among traits was also calculated from BLUP values.

### Population structure and Genome-Wide Association Analyses

Inherent population structure and cryptic relatedness can lead to spurious associations in GWAS (Astle and Balding, 2009). To control for these confounding factors, three standard GWAS models were compared: a simple one-way ANOVA model with no correction (naïve model); a general linear model (GLM) with the first five PCs of the SNP matrix as covariates (GLM + 5PCs); and a mixed-linear model (MLM) using the five PCs and marker-estimated kinship matrix (Yu et al., 2006). The models correcting for kinship and 5 PCs had the lowest inflation-factors as determined from quartile-quartile (QQ) plots and therefore the lowest false-discovery rate (**Supplementary Figure 1**). The association analyses were implemented in TASSEL (Bradbury et al., 2007; Zhang et al., 2010). Association test P-values were considered significant when more extreme than the Bonferroni threshold (with experiment-wise type I error rate of 0.05).

The patterns and extent of linkage disequilibrium (LD) in a population not only determines the obtainable resolution in association mapping studies (Hamblin et al., 2011) but also has strong implication in the interpretation of association peaks. Therefore the level of LD decay and the local patterns of LD along each chromosome were determined by calculating intra-chromosomal pairwise squared correlation (r^2^) using PLINK (Purcell et al., 2007).

## RESULTS

### SNP genotyping

A total of 72,279 genome-wide SNP markers were called for the 672 genetic gain individuals after filtering for minor allele frequency threshold of 0.005. The high-density coverage of SNPs resulted in an average of 4015 markers per chromosome, ranging from 3101 on chromosome 16 to 5880 on chromosome 1.

### Phenotypic variability

We investigated the phenotypic variation in dry matter content as well as carotenoid-based intensity of yellow root color using a visual color chart and chromameter. The dry matter content varied widely in the Genetic Gain population ranging from 8.4% to 45% (average 28.6%, Table 1). About two-thirds of the evaluated clones have white storage roots while the remaining showed a range of yellow color suggesting varying levels of carotenoid content. On average, the visual score was 1.7 and ranged from 1 (white) to 7 (yellow). The average chromameter measure of yellow color intensity (b* value) was 20.8 and ranged from 11.1 (white) to 40.8 (yellow). Dry matter was approximately normally distributed while chromameter b* values showed a bimodal distribution (**Figure 1**) in which the first peak (b* values from 10 to 20) is associated with the white clones while the second peak (b* values from 20 to 40) is associated with the variations among the yellow clones.

**Table 1.**
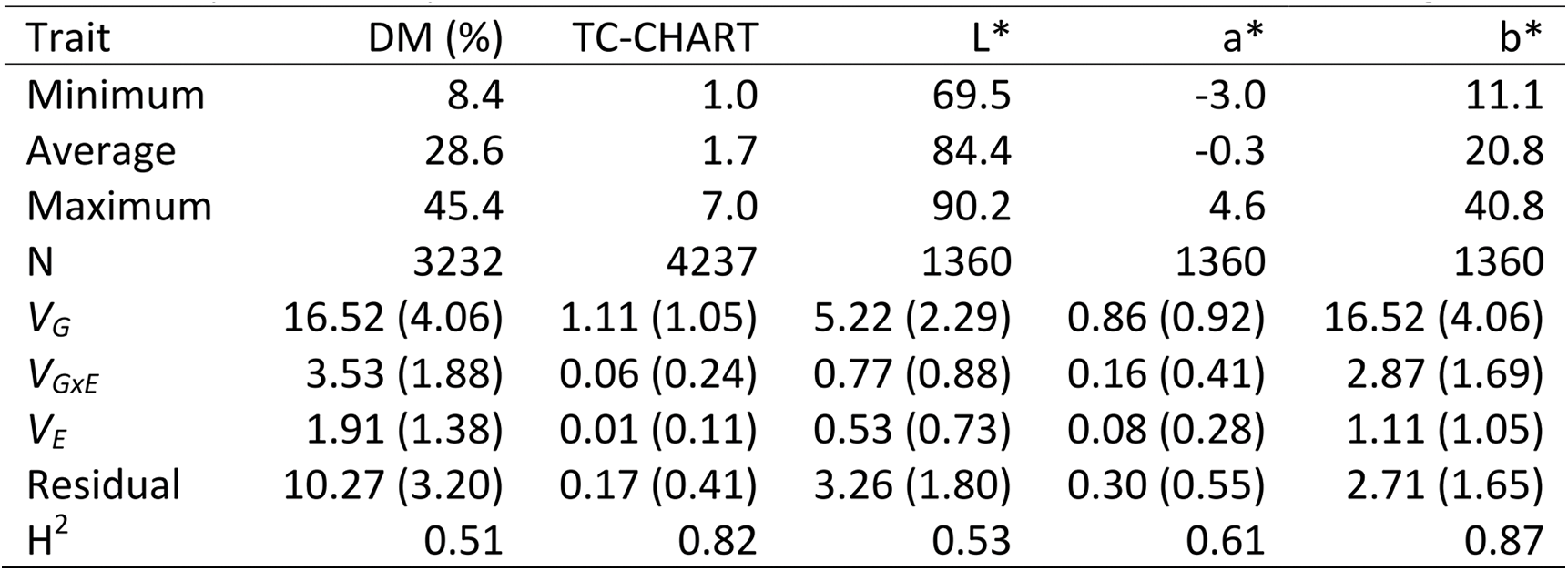
Summary of phenotype variation, variance components (±se) and broad-sense heritability (H^2^) for dry matter, color chart and Chromameter CIELAB readings.

| Trait | DM (%) | TC-CHART | L* | a* | b* |
| --- | --- | --- | --- | --- | --- |
| Minimum | 8.4 | 1.0 | 69.5 | -3.0 | 11.1 |
| Average | 28.6 | 1.7 | 84.4 | -0.3 | 20.8 |
| Maximum | 45.4 | 7.0 | 90.2 | 4.6 | 40.8 |
| N | 3232 | 4237 | 1360 | 1360 | 1360 |
| $V_G$ | 16.52 (4.06) | 1.11 (1.05) | 5.22 (2.29) | 0.86 (0.92) | 16.52 (4.06) |
| $V_{G \times E}$ | 3.53 (1.88) | 0.06 (0.24) | 0.77 (0.88) | 0.16 (0.41) | 2.87 (1.69) |
| $V_E$ | 1.91 (1.38) | 0.01 (0.11) | 0.53 (0.73) | 0.08 (0.28) | 1.11 (1.05) |
| Residual | 10.27 (3.20) | 0.17 (0.41) | 3.26 (1.80) | 0.30 (0.55) | 2.71 (1.65) |
| $H^2$ | 0.51 | 0.82 | 0.53 | 0.61 | 0.87 |

**Figure 1.**
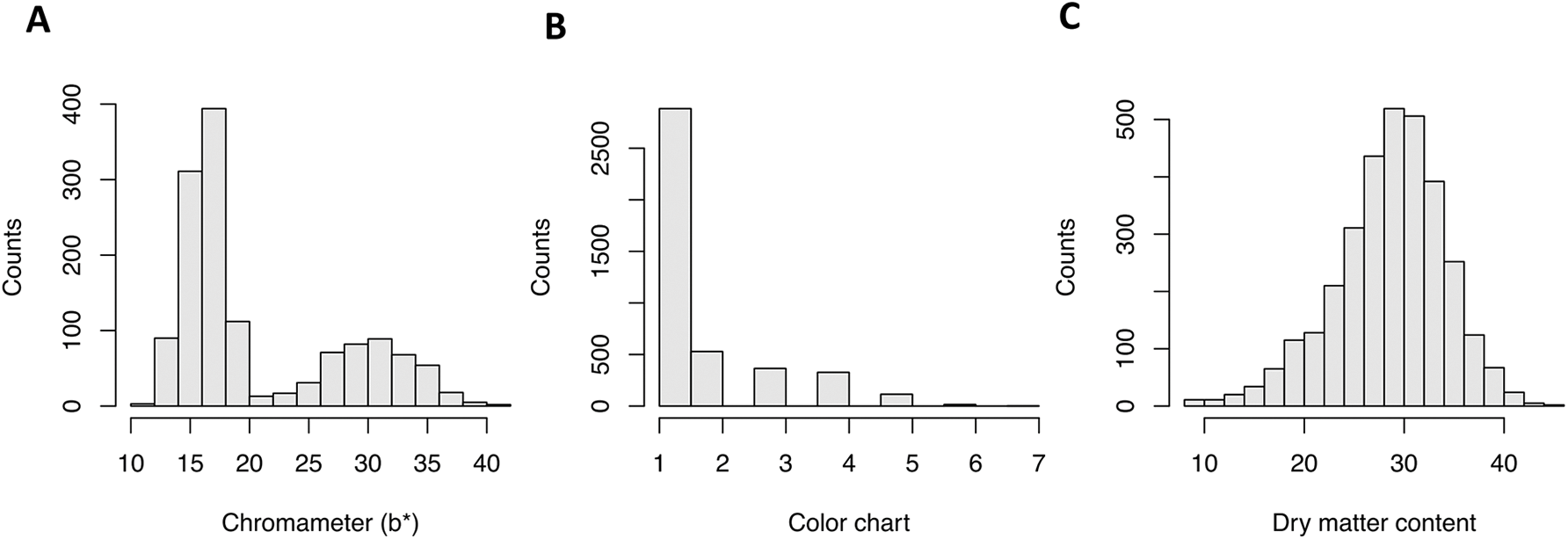
Distribution of phenotype for TCHART, Dry matter content, and chromameter (b*).

Broad-sense heritability was high for root yellow color (*H*^2^ = 0.87 and 0.82 for color chart and chromameter b* values, respectively) but moderate for dry matter content (*H*^2^ = 0.51). These values are within the range of heritabilities reported previously for these traits (Balyejusa Kizito et al., 2007; Ceballos et al., 2013). The relative importance of genotype-by-environment variance (*V*_*GxE*_) compared to genotype (*V*_*G*_) variance was measured by the ratio *V*_*GxE*_*/V*_*G*_. For all traits, the genetic variance component was larger than the genotype-by-environment interaction variance. The interaction is minimal for the yellow color measurements (0.054 and 0.173 for color chart and chromameter b* values, respectively). For dry matter content, we observed a slightly higher interaction ratio of 0.214.

The BLUPs for dry matter content and gradation of yellow color were negatively correlated in our germplasm collection (Pearson’s correlation coefficient, *r* = -0.59; P-value < 0.0001), indicating that clones with higher carotenoid content are more likely to have low dry matter content (**Figure 2**) which confirms previous findings in cassava (Marín Colorado et al., 2009; Akinwale et al., 2010; Esuma et al., 2012; Njoku et al., 2015). On the other hand, we found a positive association between dry matter content and color lightness (chromameter L* value, *r* = 0.60, P-value < 0.0001). The two measures of yellow color (i.e. color chart and chromameter b* axis) were strongly correlated (*r* = 0.96, P-value < 0.0001).

**Figure 2.**
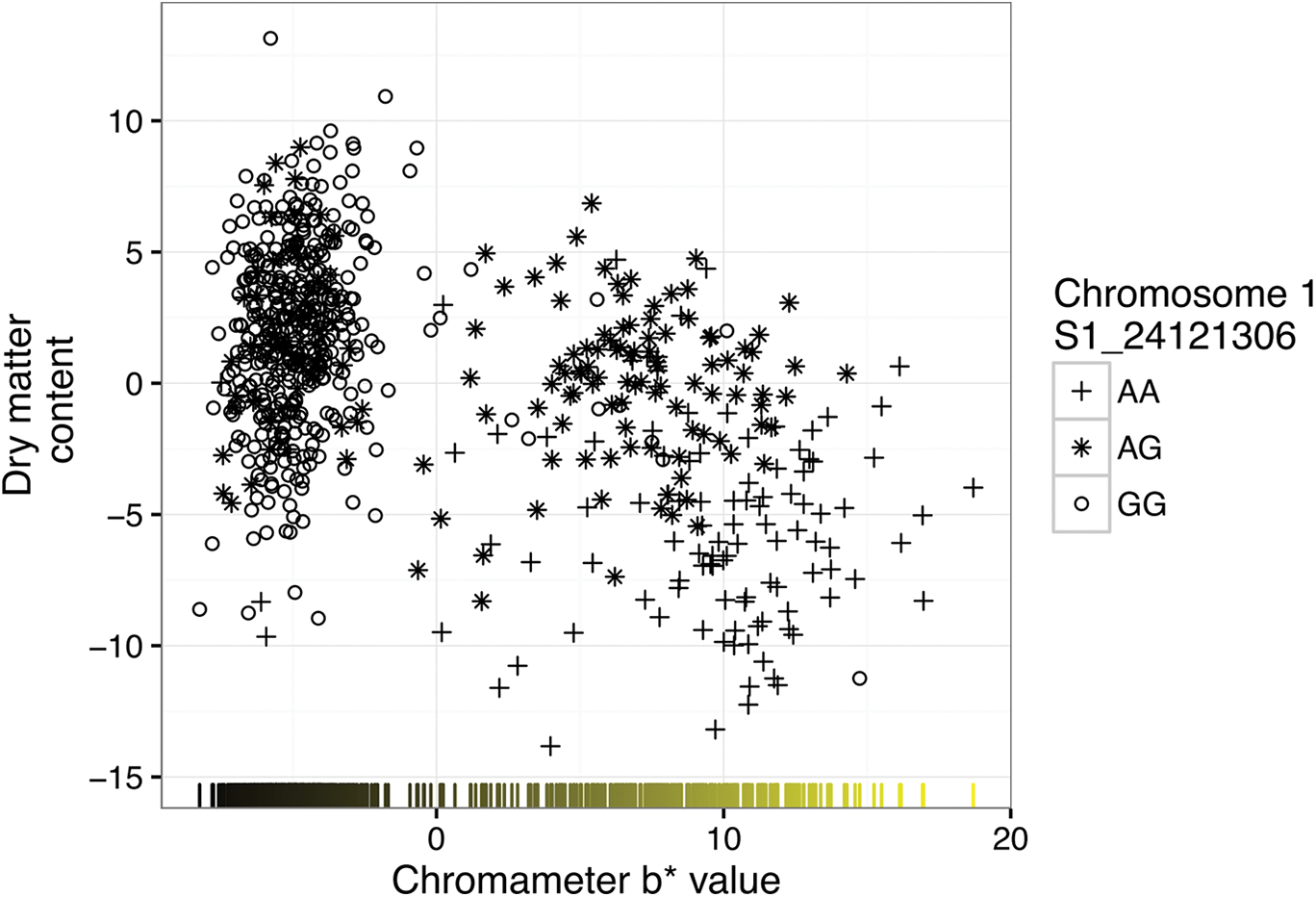
Relationship between dry matter content and root yellowness BLUPs (expressed as b* value of chromameter measurement). Different symbols denote the genotype at marker S1_24121306 that is associated with both dry matter content and root color intensity.

### Population structure

Analysis of population structure in 672 accessions genotyped across 72,279 SNPs using PCA detected subtle genetic differentiation in the genetic gain collection, with the first 10 PCs explaining about 23 % of the genetic variation. The first two principal components, which accounted for 8% of the genetic variation, revealed genetic differentiation between white and yellow-root clones (**Figure 3**).

**Figure 3.**
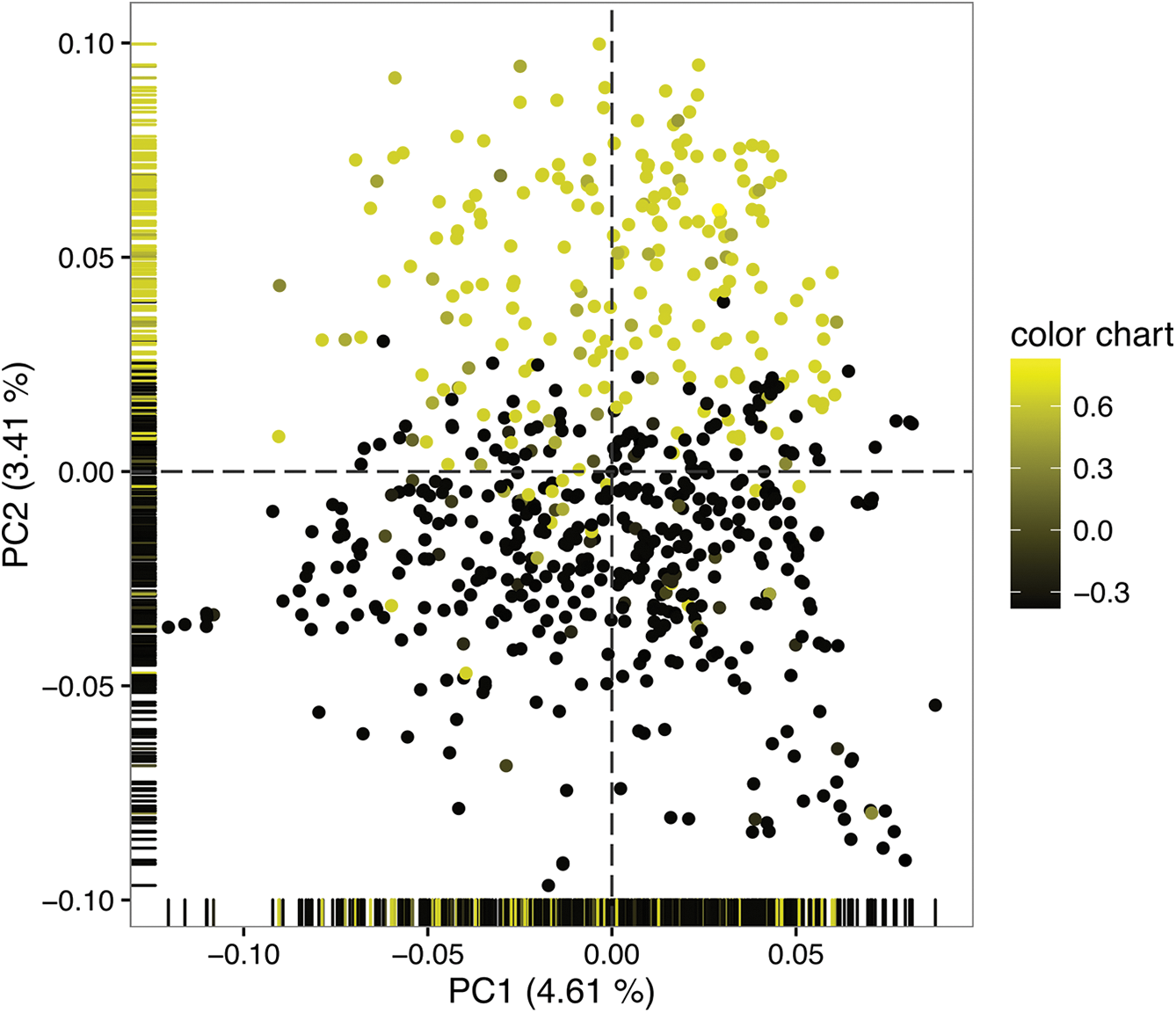
Population structure of the Genetic Gain collection. (A) PCA bi-plot of the first two axes; (B) Neighbor-joining dendrogram calculated from pairwise IBS distance. Yellow color highlights accessions with yellow roots.

### Linkage disequilibrium

Several regions of extensive mega-base-scale LD were discovered in chromosomes 1, 4 and 10 as well as smaller regions in other chromosomes (**Figure 4**).Excluding results from chromosomes with large LD blocks (i.e. chromosomes 1, 4 and 10), we found that on average, LD drops almost to background levels (r^2^ < 0.1) at around 2 Mb in the Genetic Gain population (**Figure 5**).

**Figure 4.**
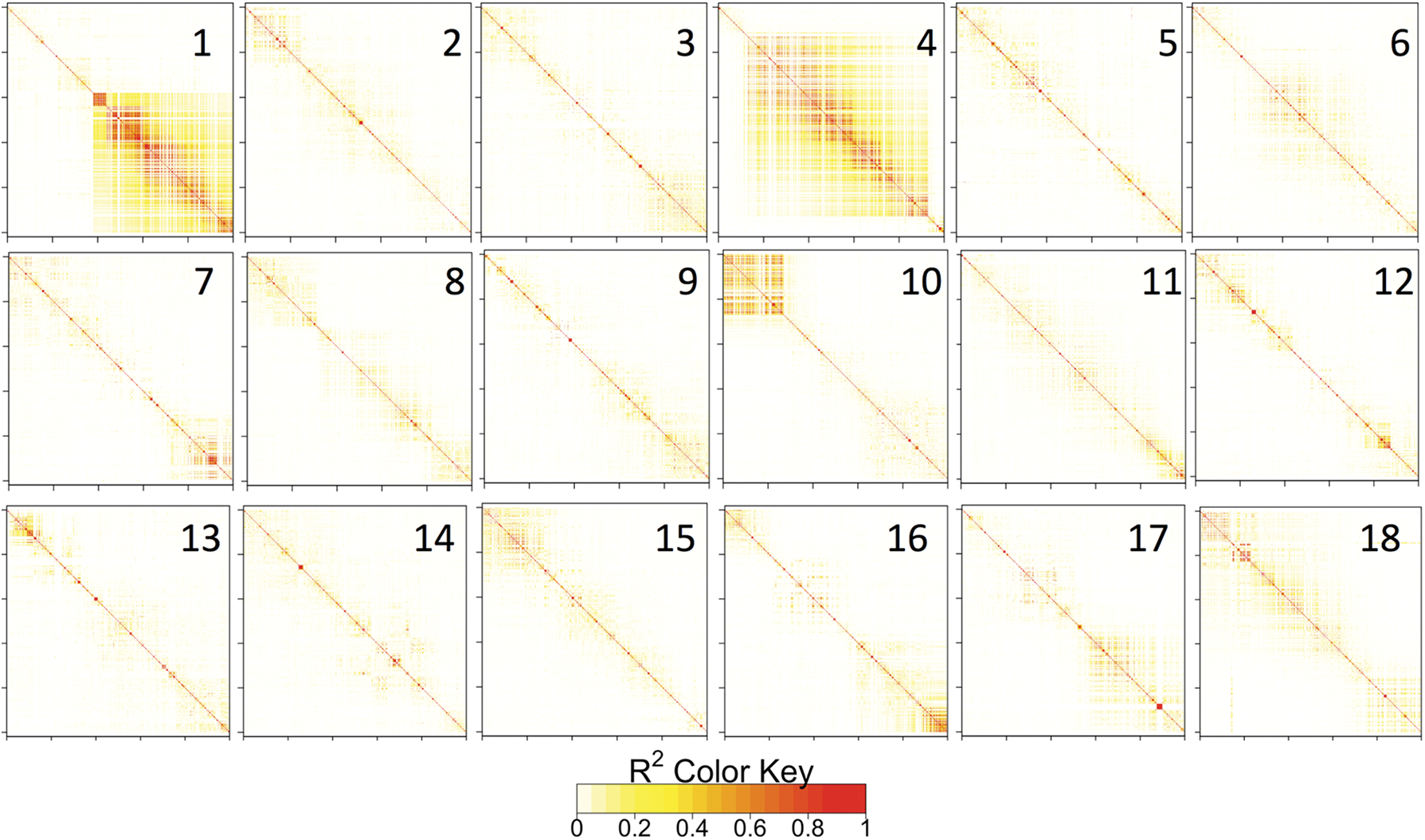
Local pattern of linkage disequilibrium (r^2^) along each of the 18 cassava chromosomes. Note the large LD blocks in chromosomes 1, 4 and 10. SNPs are arrayed according to their order, and not their physical position.

**Figure 5.**
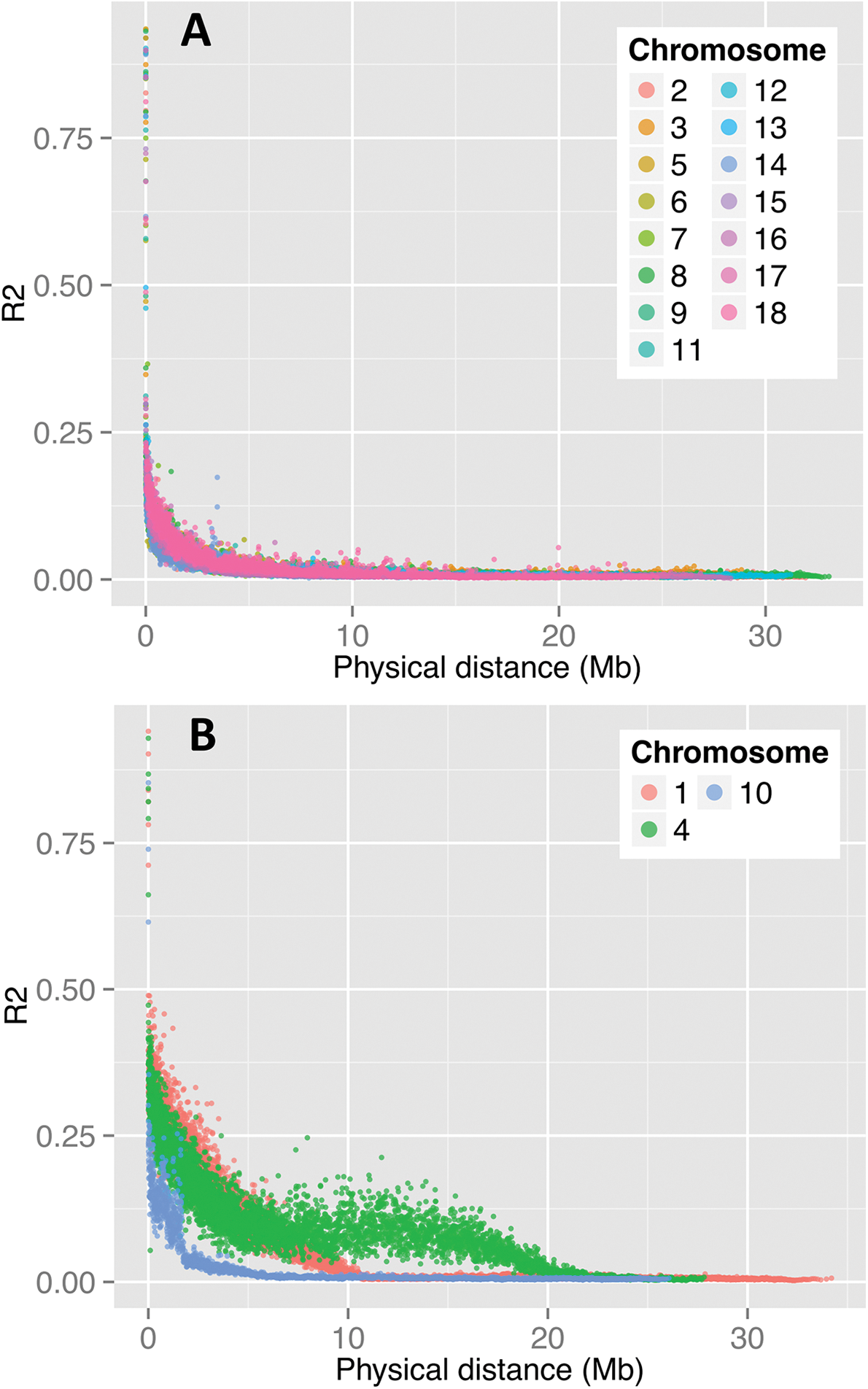
A Moving-average based LD decay profile in the Genetic Gain population. (A) all chromosomes except 1, 4 and 10; (B) chromosomes 1, 4 and 10.

## Population-wide GWAS

### Variation in carotenoid content estimated by root yellowness

The MLM-based GWAS analysis for yellow color in the storage root parenchyma using both the color chart and the chromameter-based methods uncovered the same major association regions occurring at 24.1 and 30.5 Mbp of chromosome 1 (**Figure 6**). This is not surprising given the high correlation between the two color assessment methods. The first major peak is tagged by marker S1_24121306 for visual gradation of color (-log_10_(p-value) of 21.8) and marker S1_24159585 (-log_10_(p-value) of 18.8) for chromameter b* value (Table 2). The second peak was tagged by the same marker, S1_30543382, for both measures of color intensity and occurred 6.5 Mbp away from the first peak (-log_10_(p-value) of 10.54 and 10.78, for color chart and chromameter, respectively). All other SNPs between the two major regions were not significant at the Bonferroni significance threshold (P = 6.92e-07) (**Figure 7**). The LD between SNPs S1_24121306 and S1_30543382 was 0.3, suggesting moderate non-random segregation of alleles of the two markers in these regions.

**Figure 6.**
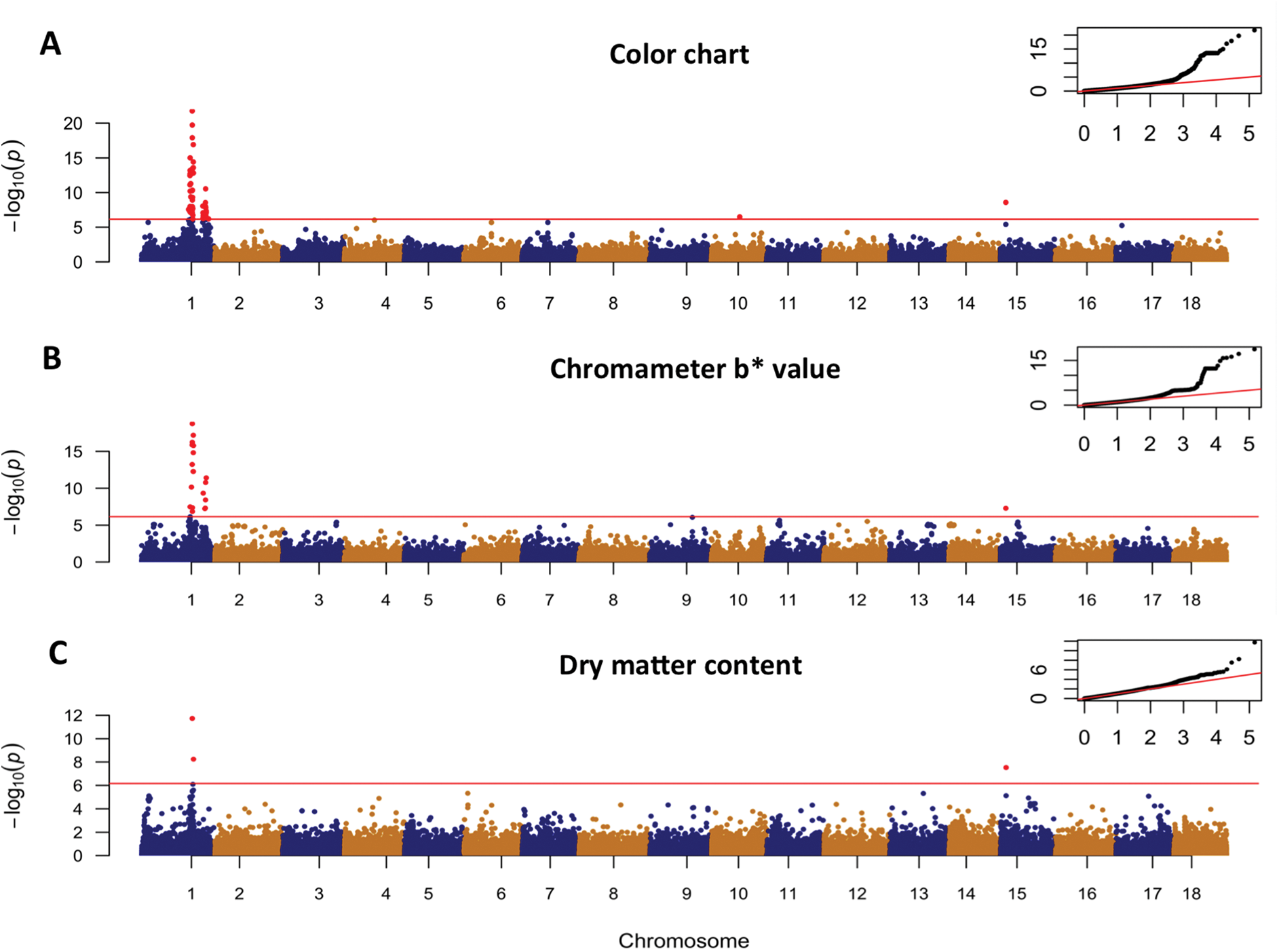
Genome-wide association results. Manhattan and Quantile-quantile plot of the MLM model for: root yellowness estimated using (A) chromameter b* value; and (B) color chart method; and (C) dry matter content. The red horizontal line indicates the genome-wide significance threshold.

**Table 2.** Summary of significant associations between selected traits and SNP markers from the MLM analysis. Only results in the major loci from Chromosome 1 are shown.

| Trait | SNP | Chr | Position (bp) | P-value | Candidate genes and mid-position (bp) |
| --- | --- | --- | --- | --- | --- |
| Color chart | S1_24121306 | 1 | 24,121,306 | 1.74E-22 | Phytoene synthase (Manes.01G124200; 24,155,070 bp) |
| Color chart | S1_30543382 | 1 | 30,543,382 | 2.91E-11 | NA |
| Chromameter b* | S1_24159585 | 1 | 24,159,585 | 1.79E-19 | Phytoene synthase (Manes.01G124200; 24,155,070 bp) |
| Chromameter b* | S1_30543382 | 1 | 30,543,382 | 1.66E-11 | NA |
| Dry matter | S1_24121306 | 1 | 24,121,306 | 1.86E-12 | UDP-glucose pyrophosphorylase (Manes.01G123000; 24,061,652 bp); sucrose synthase (Manes.01G123800; 24,142,314 bp) |

Chr = Chromosome (version 6 of cassava reference genome);

**Figure 7.**
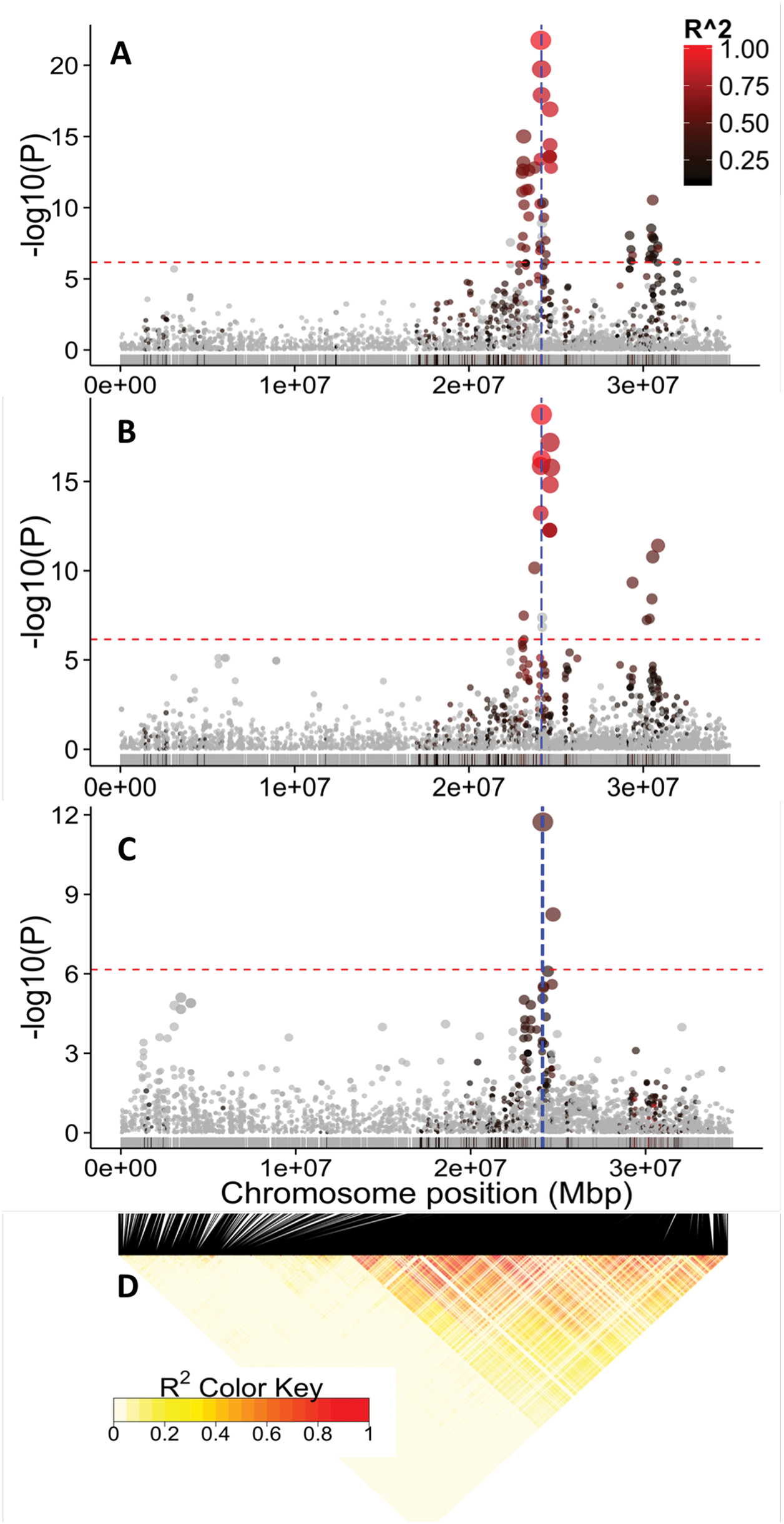
GWAS results for chromosome 1. Manhattan and Quantile-quantile plot of the MLM model for: (A) root yellowness as measured by chromameter b* value; (B) color chart; and (C) dry matter content. Note the common peak at ~ 24.1 Mbp region for the three traits. Red horizontal line indicates the genome-wide significance threshold. The SNPs are colored according to their degree of linkage disequilibrium (r^2^) with the leading variant (i.e. top SNP for the first peak at 24.1 Mbp). The vertical blue lines in (A) and (B) denote the position of the carotenoid biosynthesis gene, phytoene synthase (24,155,070 bp), and those on (C) denotes the positions of the UDP-glucose pyrophosphorylase (24,061,652 bp) and sucrose synthase (24,142,314 bp).

### Dry matter content

Genetic variation in dry matter content was found to be associated with a major locus occurring at 24.1 Mbp region of chromosome 1 and tagged by marker S1_24121306 (–log10(p-value) of 11.73). Importantly, this locus for dry matter content co-locates with one of the two peaks found to be associated with carotenoid content (**Figure 7**). The genomic co-location of the major loci for dry matter content and root yellowness suggests either a strong physical linkage between the genes underlying these important traits or a pleiotropic effect. Distinguishing between these two possible causes is important in cassava improvement efforts that target both traits. We therefore attempted to unravel the genetic cause of the observed association by: (i) exploring the underlying linkage disequilibrium patterns in the QTL regions on chromosome 1; (ii) carrying out independent association analysis for dry matter content within the white root and yellow root subpopulations; and (iii) searching for plausible biological explanation by identifying candidate genes for both traits in the target region.

Exploration of the LD landscape along Chromosome 1 uncovered a mega-base-scale region of low recombination extending from 22Mb to 33 Mb surrounding the association peaks for dry matter content and yellow color (**Figure 7**). This region was recently shown to coincide with a large *Manihot glaziovii* introgression segment (Bredeson et al., 2016) that traces back to early breeding for resistance to cassava mosaic and cassava brown streak viruses in the 1930’s (Hahn et al., 1980). Clustering of the Genetic Gain population based on identity-by-descent relationship (i.e. a measure of how many alleles at any marker in each of the two samples came from the same ancestral chromosomes) calculated using only markers from this extensive LD region (2150 SNPs from markers S1_21567540 to S1_34950326) revealed at least two major groups of accessions (**Supplementary Figure 2**), indicating presence of few major haplotypes associated with the LD blocks.

### GWAS for dry matter content in white root and yellow root subpopulation

If the phenotypic association between dry matter and carotenoid contents and the colocation of their association signals (~ 24.1 Mbp region) is largely caused by physical linkage rather than by pleiotropy, the major dry matter locus should be detectable in both white root and yellow root germplasm when analyzed independently. We therefore split the Genetic Gain dataset into white root (n=210) and yellow root (n=427) subpopulations and repeated the GWAS analysis. Clones that were at the borderline between yellow-root and white-root were excluded from these analyses. To mitigate the loss of power as a result of double-fitting markers in the MLM model both as a fixed effect tested for association and as a random effect as part of the kinship (Lippert et al., 2011; Listgarten et al., 2012), the MLM analysis was carried out using a kinship matrix calculated excluding markers from chromosome 1.

We recovered the major dry matter content association signal in both the white root and yellow-root subpopulation (**Figure 8**). Though coinciding with the locus identified in the population-wide GWAS, the association signal in the white subpopulation was much broader, extending from 24 to 33 Mbp and generally overlaps with the broad LD region of the chromosome 1 (**Figure 8**). On the contrary, association signal for the yellow subpopulation was relatively narrow. Survey of the underlying LD pattern in the same chromosome region for the yellow subpopulation showed a recombination spot.

**Figure 8.**
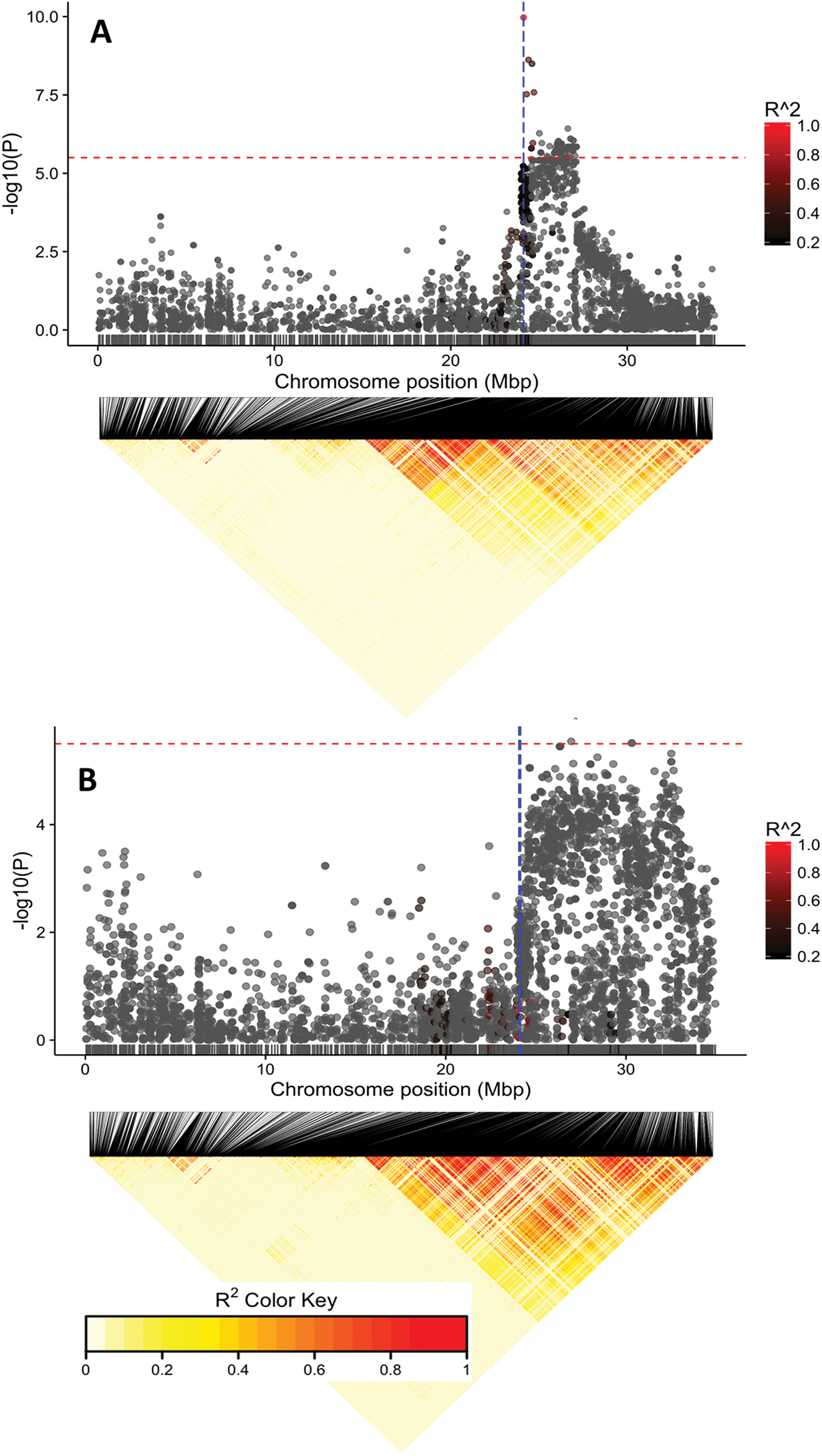
Manhattan plots of the MLM analysis of the yellow root (top) and white root subpopulations (bottom). Below each is an LD heatmap showing pairwise squared correlation of alleles between markers along chromosome 1. Note the large number of SNPs showing significant association with dry matter in the white subpopulation compared to that of the yellow subpopulation. Red horizontal line indicates the genome-wide significance threshold. The vertical blue lines are same as in Figure 7.

### Selection sweep associated with breeding for yellow-root varieties

To determine whether the breeding for carotenoid content trait in the Genetic Gain germplasm resulted in a selection sweep around the major QTL region, we quantified genome-wide nucleotide variation in the yellow root subpopulation (n = 210) and the non-yellow subpopulation (n = 427). A sliding-window scan of expected heterozygosity (π) and Tajima’s D detected a ~ 6 Mb region with decrease in nucleotide diversity in the yellow compared to white-root subpopulation around the first major carotenoid locus site (~ 24.1 Mbp) relative to its chromosomal neighborhood (**Figure 9**).

**Figure 9.**
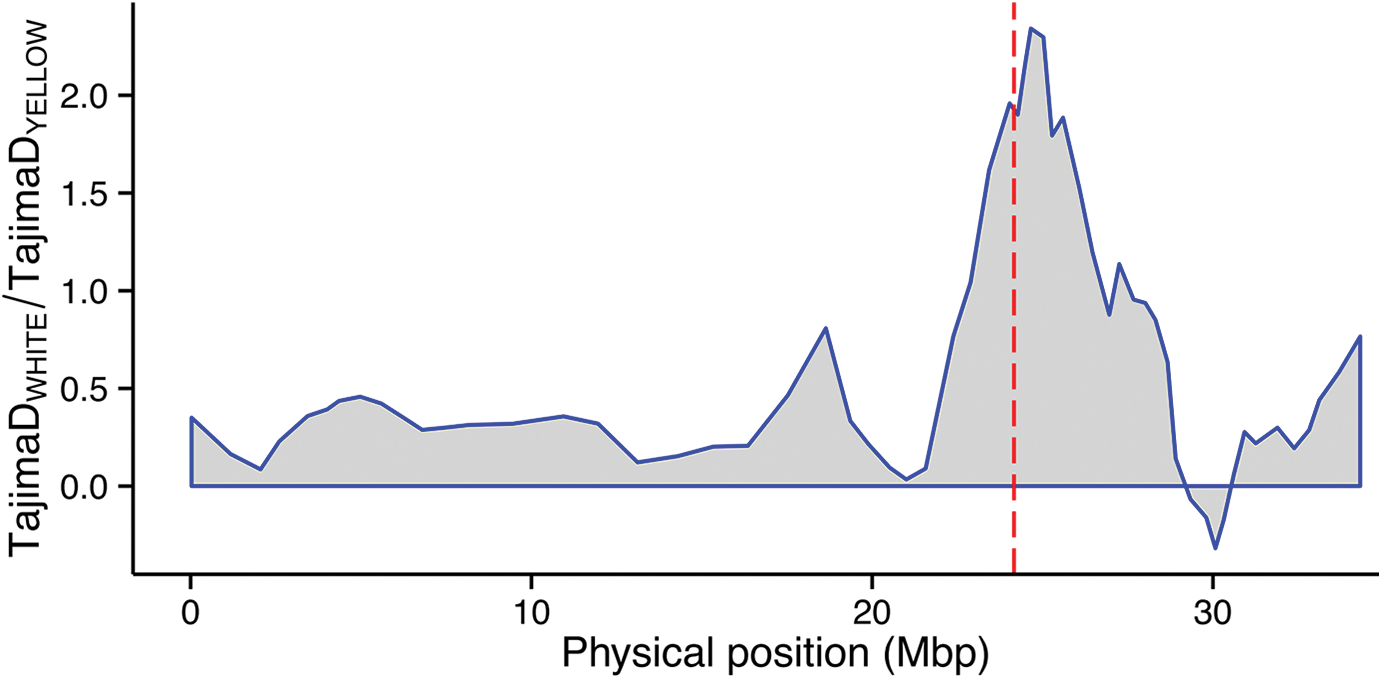
Selection sweep associated with positive selection for provitamin A trait varieties in the genetic gain population. Red dashed line indicates the position of SNP S1_24121306.

### Proportion of variance explained by markers QTL haplotypes

To determine predictive ability of the discovered loci for yellow color intensity and dry matter content, we carried out a multiple linear regression analysis using the *lm* function in R and considered the top markers for these traits as independent variables and the traits measurements as the response variables. A model considering the two major peaks associated with gradation of yellow color as assessed using a color chart (S1_24121306 and S1_30543382) returned an adjusted squared correlation (R^2^) of 0.81. For the measure of continuous variation in intensity of yellow color using chromameter (b* value), the adjusted R^2^ from same genomic regions (S1_24159585 and S1_30543382) was 0.70 while that for dry matter content was moderate (R^2^ = 0.37). This finding suggests that the major loci on Chromosome 1 would be useful in Marker Assisted Selection breeding in cassava. Single or joint allelic substitution effects at the associated loci with respect to chromameter b* value, color chart and dry matter content is shown in **Figure 10**.

**Figure 10.**
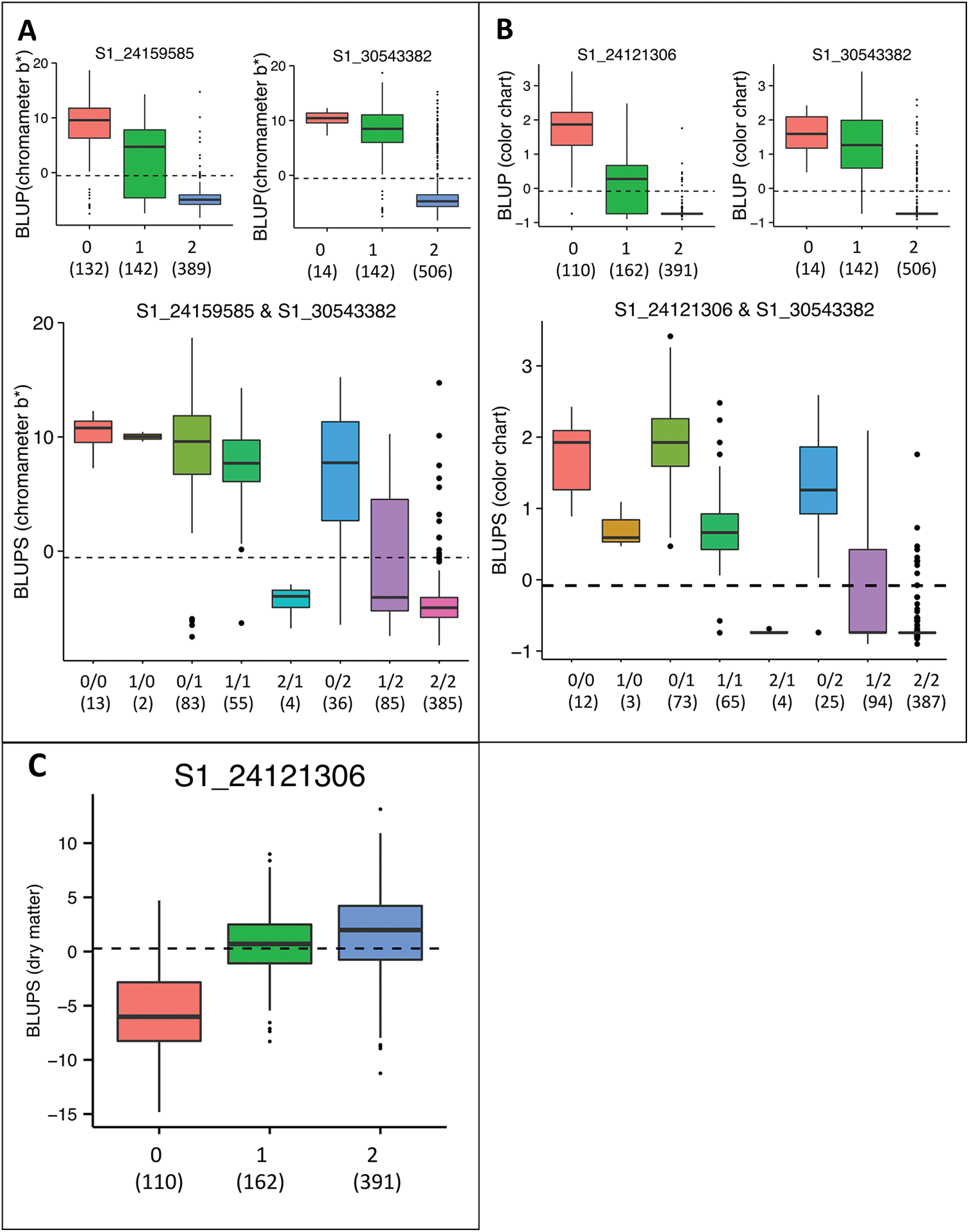
Effect of the most significantly associated markers on the BLUPs for yellow color measured by: (A) Chromameter (b* value) and (B) color chart in the Genetic Gain population. The boxplots show the effects of the most significantly associated SNPs at first and second peaks (above) and the two-locus haplotypes (below) on chromosomes 1. (C) Effect of the most significantly associated markers on the BLUPs for dry matter content. Alleles are coded as 0 = homozygous reference genome; 1 = heterozygous and 2 = homozygous non-reference genome. The dashed line represents the population mean of the BLUPs. The numbers in parenthesis below genotypic categories refer to the number of accessions for each genotype.

### Candidate genes

The first of the two genomic regions associated color intensity (tagged by SNP S1_24159585) was found ~ 4.5 Kbp away from phytoene synthase 2 (*PSY*2) in the cassava version 6 reference genome. The *PSY*2 enzyme, named Manes.01G124200 and located at 24,155,070 bp, is involved in the first dedicated step of the carotenoid biosynthesis pathway in cassava roots which converts geranylgeranyl diphosphate to phytoene (Welsch et al., 2010). Presence of the null versus the functional *PSY*2 allele is responsible for the qualitative color difference between the white and the yellow roots, respectively (Welsch et al., 2010; Rabbi et al., 2014a). Our study suggests that allelic variation associated with increases in enzyme activity could contribute to deeper yellow by increasing the flux into the pathway. No known candidate genes were found in the vicinity of the second significant association signal on chromosome 1 occurring at 30.5 Mbp.

For dry matter content, we found two particular genes that are pivotal in central carbon metabolism in the vicinity of top SNP linked to that trait. The first is UDP-glucose pyrophosphorylase (named Manes.01G123000 in the cassava reference genome). This gene which occurs at 24.06 Mbp region, plays a key role in carbohydrate metabolism, and is strongly associated with the yield production both in grains and root crops (Smith, 2008; Zeeman et al., 2010). UDP-glucose pyrophosphorylase was recently found to be up-regulated during bulking of cassava storage roots (Yang et al., 2011; Wang et al., 2016). The second key carbohydrate metabolism gene was sucrose synthase (named Manes.01G123800), which occurred in 24.14 Mbp region. Finding of these potential candidate genes for carotenoid and carbohydrate biosynthesis strongly favors the possibility that the association between these two traits is caused by physical linkage rather than pleiotropy. This hypothesis warrants further investigation.

## DISCUSSION AND CONCLUSION

The present study revealed that the genetic architecture for dry matter content and intensity of yellow color resulting from carotenoid accumulation in cassava roots is governed by few major loci on chromosome 1 and explains the large repeatability estimates, particularly for yellow color. These findings expand on those from previous genetic mapping efforts for dry matter and carotenoid content. Using a candidate gene mapping approach, Welsch et al. (2010) reported that a SNP mutation in the *PSY2* gene, leading to amino-acid substitution, differentiates white and yellow cassava storage roots. Similarly, a bi-parental QTL mapping study that used two clones from the Genetic Gain collection (TMS-I961089A and TMEB117) also uncovered a single QTL peak whose confidence interval encompassed the same PSY2 gene (Rabbi et al., 2014a). The F1 progenies from the TMS-I961089A x TMEB117 population, also genotyped using the GBS method, segregated at an approximately 1:1 ratio for white versus light-yellow roots, suggesting that the yellow-root parent was heterozygous for the functional allele at the PSY2 locus. Kizito et al. (2007) reported a QTL for dry matter content in a bi-parental population genotyped using SSR markers that also corresponds to this region on chromosome 1. More recently, Esuma et al., (2016) reported a single genomic region on Chromosome 1 underlies the variation in total carotenoid content in eight S1 and S2 partially inbred families. This peak, around 24.66 Mbp, is close to our first locus tagged by SNP S1_24121306. However that study did not look at genetic architecture for dry matter content.

While the amount of total carotenoids in the Genetic Gain collection was not directly estimated, previous studies of diverse cassava germplasm have consistently reported a strong linear relationship between yellow color and carotenoid content (Pearson’s coefficient, *r*, ranging from 0.81 to 0.89) (Iglesias et al., 1997; Chávez et al., 2005; Marín Colorado et al., 2009; Akinwale et al., 2010; Sánchez et al., 2014). Hence the results obtained here should be useful for breeding efforts targeting breeding for improved carotenoid content. Nevertheless, we propose to quantify total carotenoids and its constituents as a future study to corroborate the current findings.

Given the importance of dry matter content in cassava, and the fact that we found a single genomic region associated with this trait, further studies are warranted to fine-map and validate the identity of the causal locus. To do this effectively would require different populations that are lacking the wild introgression segments in chromosome 1. This will lead to reduced LD and allow higher mapping resolution. Additionally, special crosses such as nested-association mapping population design (Yu et al., 2008) using strategically selected sets of parents will reduce the confounding effect of population structure. Given our marker density and sample size, this study is sufficiently powered to find large effect alleles that are common in the studied germplasm. To detect more QTLs of small effects will require a larger association panel genotyped at higher density.

The use of a broad cassava diversity panel in GWAS not only provides the foundation to map genomic regions associated with natural variation in dry matter and carotenoid content but also allows us to unravel the genetic cause of the negative correlation between these traits, that is, pleiotropy versus genetic linkage. In the context of breeding to simultaneously increase carotenoid and dry matter content, the observed negative association between these traits in our germplasm is undesirable. Several lines of investigation pointed to a possibility of genetic linkage rather than pleiotropy to be the cause of the observed association. Firstly, the genomic region harboring the QTLs for yellow color and dry matter content was found to occur in chromosomal segments that exhibits low overall recombination in this region compared to the genome-wide patterns. Recent work by Bredeson et al. (2016) has shown that this chromosome 1 region harbors a large *M. glaziovii* introgression that commonly occurs in the Genetic Gain collection. Secondly, independent association analysis for dry matter content on the white and the yellow subpopulations detected the same association signal although the QTL in the white subpopulation was broader suggesting that the favorable alleles were located in non-recombining haplotype. Thirdly, strong candidate genes for dry matter (UDP-glucose pyrophosphorylase and sucrose synthase) and carotenoid content (phytoene synthase) were found in the vicinity of the major association region (24.1 Mbp) of chromosome 1. Presence of these genes hints at possibly distinct biological causes of the observed associations with the two traits. These hypotheses need to be tested through functional genetics studies at these candidate genes. Taken together, these findings suggest that the phenotypic correlation between dry matter and carotenoid content is mainly caused by physical linkage of loci underlying these trails. Moreover, Ortiz et al. (2011) found a fairly large positive correlation (r = 0.62) between these traits. It is therefore possible that the nature of association (whether positive or negative) is dependent on the allelic status at the linked dry matter and carotenoid biosynthesis genes. We also detected a reduction of expected heterozygosity (π) around the major gene region in the yellow versus white sub population. This suggests that the genetic base for sources of favorable alleles with respect to carotenoid biosynthesis at this locus is narrow, possibly arising from a single haplotype, which could be linked in *cis* to low-dry matter alleles in the dry matter locus. Alternatively, balancing selection of the *M. glaziovii* introgression in the white cassava sub population might be the cause of the higher levels of heterozygosity relative to the yellow sub population.

Although cassava is a predominantly outcrossing species, its clonally propagated nature means that modern varieties have undergone relatively few recombination cycles compared to seed crops. Most accessions in the Genetic Gain collection are not far removed from founder clones. Accordingly, the extent of LD in this study (~ 2 Mbp) is much greater than the LD in maize (< 10 Kb) (Yan et al., 2009) as well as in grape (< 10 Kb) (Myles et al., 2011), another clonal species. Moreover, the overall recombination pattern is far from homogeneous owing to the persistent introgressions of *M. glaziovii* chromosomal segments that are legacies of the historical breeding program in East Africa (Hahn et al., 1980; Jennings, 1994). From these results, it is expected that the mapping resolution will vary widely across the cassava genome depending mainly on whether a locus-of-interest occurs in or outside the large-LD blocks.

This study presents a significant progress toward dissecting the genetic architecture of two key breeding goal traits in cassava. The major loci associated with carotenoid content variation and a single locus associated with dry matter content represents markers that will be useful for marker-assisted selection in these traits. Although the results of the present study suggests genetic linkage is more likely to be responsible for the negative correlation between the studied traits, there is need for further investigations to confirm or reject this hypothesis. For example, will dry matter content be increased by knocking out the PSY2 gene using gene-silencing methods (Lu et al., 2003; Burch-Smith et al., 2004; Fofana et al., 2004)?. Alternately, could the activation of PSY2 in clones with high dry matter content and lacking in carotenoids using gene-editing technologies like CRISPR-CAS9 (Hsu et al., 2014; Sander and Joung, 2014) lead to not only carotenoid production and accumulation but also lowering of dry matter content?

## Acknowledgements

Oluwafemi Alaba and Ruth Uwugiaren for DNA processing; Staff of the Cassava Breeding Unit of IITA for conducting field trials; Sharon Mitchell, Charlotte Acharya of Cornell’s Genomic Diversity Facility; Lukas Mueller and Guillaume Bauchet for SNP processing and imputation. This work was supported through HarvestPlus project, The CGIAR Research Programme on Roots, Tubers, and Bananas (CRP-RTB) and The Next Generation Cassava Breeding grant OPP1048542 from Bill and Melinda Gates Foundation and the United Kingdom Department for International Development.

---

**Supplementary Figure 1.**
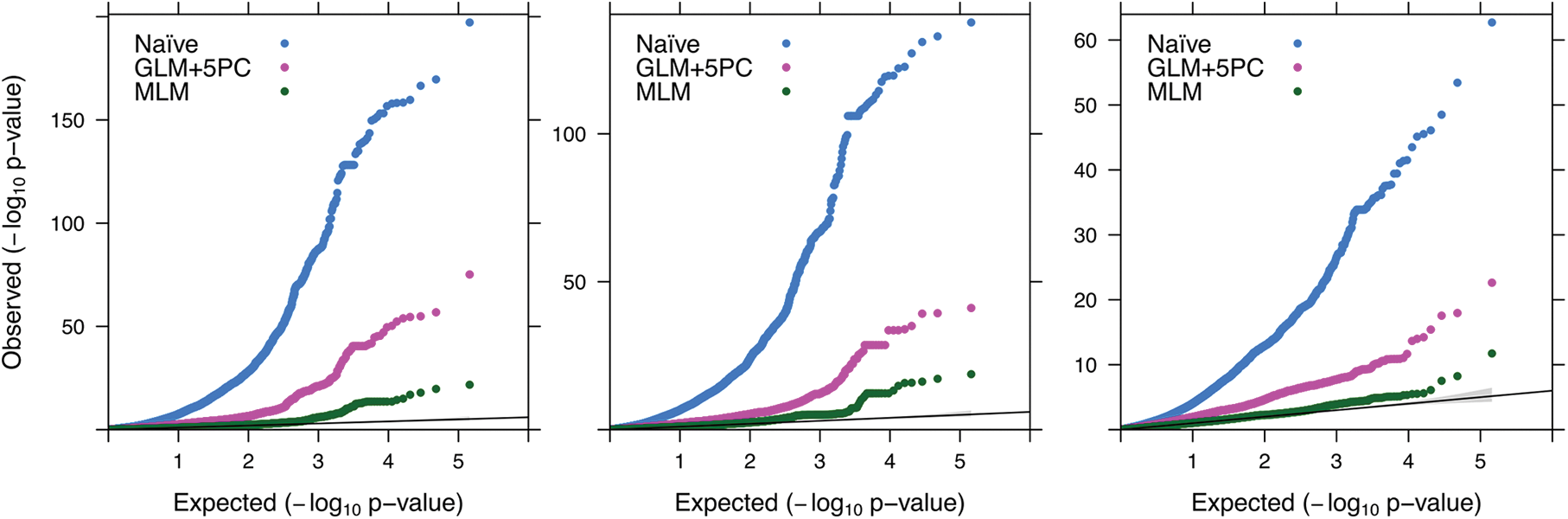
Quantile–quantile plots for P-values obtained from simple GLM, GLM+5PCs and MLM model for color chart, chromameter b* value and dry matter content.

**Supplementary Figure 2.**
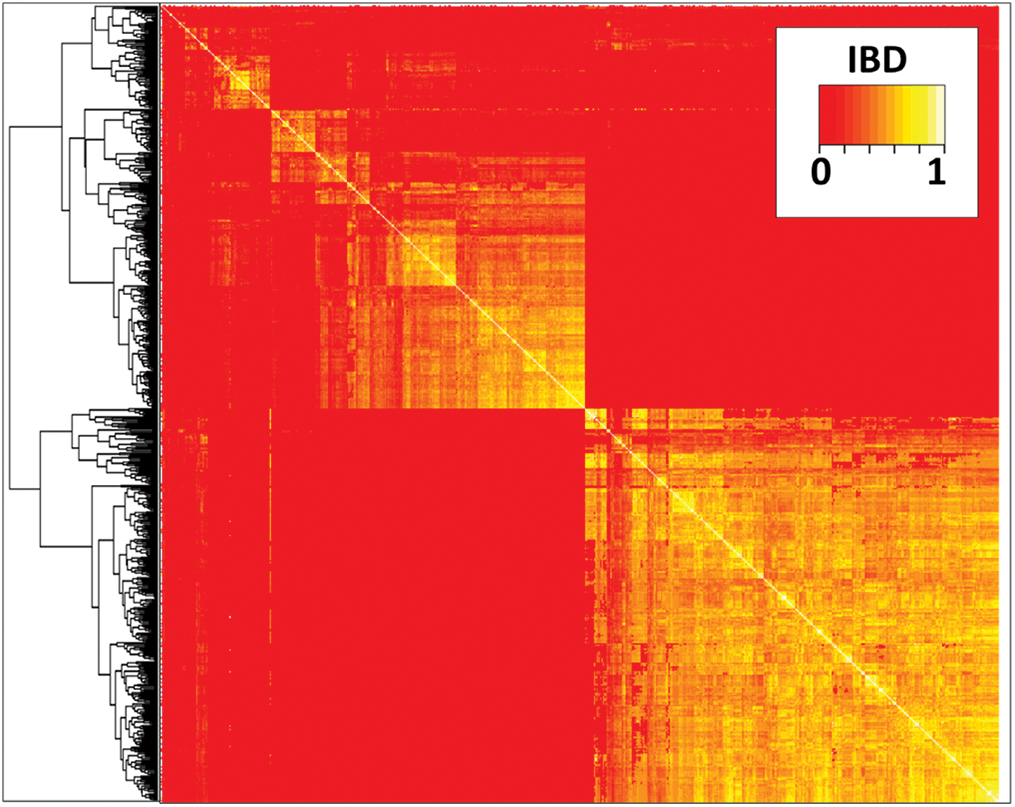
Heatmap of identity-by-descent relationship using SNPs from large LD block in chromosome 1 around the major QTL region.

